# Identification of a Zika NS2B-NS3pro pocket susceptible to allosteric inhibition by small molecules including qucertin rich in edible plants

**DOI:** 10.1101/078543

**Authors:** Liangzhong Lim, Amrita Roy, Jianxing Song

**Affiliations:** Department of Biological Sciences, Faculty of Science, National University of Singapore; 10 Kent Ridge Crescent, Singapore 119260

**Keywords:** Zika virus, NS2B-NS3 protease, Allosteric inhibition, Quercetin, Flavonoid, pNGB, Molecular docking

## Abstract

It has been recently estimated that one-third of the world population will be infected by Zika virus, but unfortunately so far there is no vaccine or medicine available. In particular, the special concern on the vaccine treatment to Zika and Dengue arising from antibody-dependent enhancement strongly emphasizes the irreplaceable role of its NS2B-NS3 protease (NS2B-NS3pro) as a target for anti-Zika drug discovery/design due to its absolutely-essential role in viral replication. Very recently we identified two small molecules inhibit Zika NS2B-NS3pro in non-competitive mode, with *K*_i_ values of 0.57 and 2.02 µM respective for p-Nitrophenyl-p-guanidino benzoate and qucertin. Here, by molecular docking, we show that although one is designed compound while another is a natural product, both molecules bind to the same pocket on the back of the substrate-binding pocket of Zika NS2B-NS3pro. As the two inhibitors fundamentally differ from cn-716, the only known peptidomimetic boronic acid inhibitor in both structure scaffolds and inhibitory modes, our discovery might open up a new avenue for the future development of allosteric inhibitors, which is highly demanded to achieve therapeutic inhibition of flaviviral NS2B-NS3pro complexes. Furthermore, as qucertin is abundant in many vegetables and fruits such caper, lovage, tea and red onion, our results should benefit the public to immediately fight Zika virus.

## INTRODUCTION

Zika virus was originally isolated from a sentinel rhesus monkey in the Zika Forest of Uganda in 1947 (1), which is transmitted to humans by Aedes species mosquitoes. Since 2007, large epidemics of Asian genotype Zika virus have been reported around the world (2–4). Recently it has been estimated that one-third of the world population will be infected (5). Most seriously, Zika infection has been found to be associated with serious sequelae such as Guillain-Barré syndrome, and microcephaly in newborn infants of mothers infected with Zika virus during pregnancy (6–9), and consequently WHO has declared a public health emergency for Zika virus (10). Zika virus represents a significant challenge to the public health of the whole world but unfortunately there is no effective vaccine or other therapy available so far.

Zika virus with a single stranded, positive sense RNA genome of 10.7 kb belongs to the flavivirus genus, which also contains Dengue, yellow fever, West Nile, Japanese encephalitis, and tick-borne encephalitis viruses (4,11). Zika virus shares a high degree of sequence and structural homology with other flaviviruses particularly Dengue virus, thus resulting in immunological cross-reactivity (7). As such, Zika was proposed as the fifth member of the Dengue serocomplex. Seriously, the current Zika outbreaks are largely localized within dengue-endemic areas, it is thus possible that preexisting dengue-induced antibodies may enhance Zika infection by antibody-dependent enhancement (ADE), a factor that makes the vaccine approaches extremely challenging (7).

Zika genome is translated into a single ~3,500-residue polyprotein, which is further cleaved into 3 structural proteins and 7 non-structural proteins (11). The correct processing of the polyprotein is essential for replication of all flaviviruses, which requires both host proteases and a viral NS2B-NS3 protease (NS2B-NS3pro) (11–18). As a consequence, the flaviviral NS2B-NS3pro has been well-established to be key targets for developing antiviral drugs (11–18). In particular, the unique concern on the vaccine treatment to Zika and Dengue strongly emphasizes the irreplaceable role of the Zika protease as a target for antiviral drug discovery/design. As facilitated by our previous studies on Dengue NS2B-NS3pro (12), we started to work on Zika NS2B-NS3pro immediately after the outbreak of Zika and successfully obtained several active forms of recombinant Zika NS2B-NS3pro. Furthermore, we attempted to discover its inhibitors from both synthetic compound library and edible plants/traditional herbal medicines to fight Zika.

## RESULTS

Recently we identified two small molecules which could amazingly inhibit Zika NS2B-NS3pro in non-competitive mode (19–21): one is p-Nitrophenyl-p-guanidino benzoate (pNGB) (Fig. 1A), an active site inhibitor for both West Nile and Dengue protease (19), while another is qucertin (Fig. 1B), a natural product rich in many edible plant (Fig. 1C).

**Fig 1.**
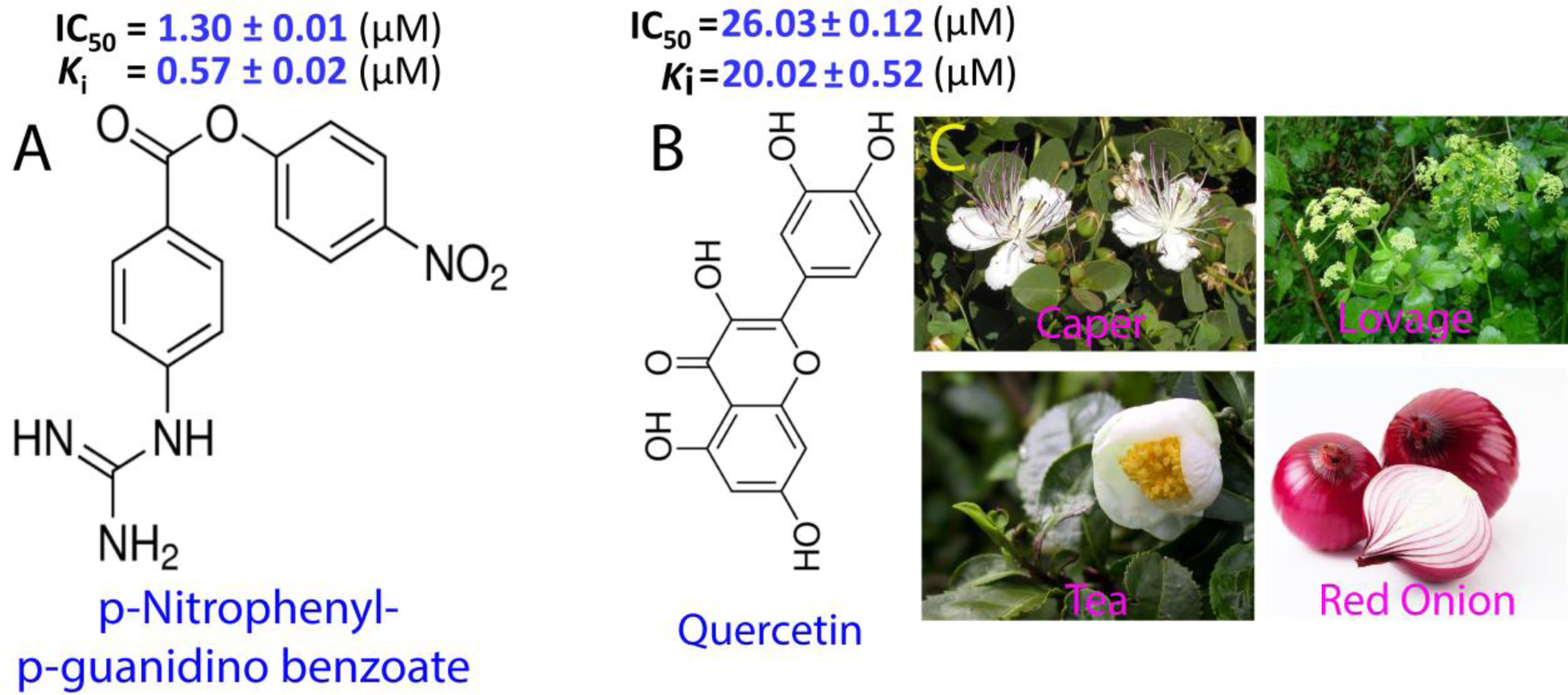
Two small molecules inhibiting Zika NS2B-NS3pro in non-competitive mode. (A) Chemical structure of p-Nitrophenyl-p-guanidino benzoate. (B) Chemical structure of Qucertin. (3) Several edible plants rich in Qucertin. The inhibitory parameters including IC_50_ and *>K*i on Zika NS2B-NS3pro are displayed for both molecules.

The results that both of them inhibit Zika protease in non-competitive mode suggest that they are allosteric inhibitors, whose binding pockets should have no overlap with the substrate-binding sites. To insight into the structural details of possible binding pockets, we decided to dock two small molecules to Zika NS2B-NS3pro. The chemical structure of quercetin (ZINC Num: ZINC03869685) and pNGB (CAS Num: 19135-17-2) were downloaded from ZINC (http://zinc.docking.org) and ChemicalBook database (http://www.chemicalbook.com) respectively. Subsequently the chemical structures were then geometrically optimized with Avogadro [22]. The partial charges of all atoms in small compounds and Zika NS2B-NS3pro were assigned with Gasteiger-Marsili charges, and non-polar hydrogen atoms were merged into the appropriate heavy atoms with AutoDockTools [23]. AutoDock software (Version 4.2) was utilized to dock Quercetin and pNGB to open conformation of Zika NS2B-NS3pro. The grid box was set with 74 × 70 × 66 (x,y,z axis) with the default 0.375Å spacing. The initial population size was set to 300, and the number of energy evaluations was set to 25,000,000, and number of docking runs was set as 150. The results were clustered with each cluster having a tolerance of 2Å. The complexes with the lowest energy were selected for analysis and display.

Fig. 2 presents the Zika NS2B-NS3pro in complex with pNGB. Indeed, as shown in Fig. 2A and 2B, pNGB binds to a pocket on the back of the substrate-binding pocket which is occupied by cn-716, a very tight competitive inhibitor for Zika NS2B-NS3pro [24]. Furthermore, as seen in Fig. 2C, the pocket is located in between two subdomains of the chymotrypsin fold, with a hydrophobic pocket at the bottom, and the rest of surface negatively charged.

**Fig 2.**
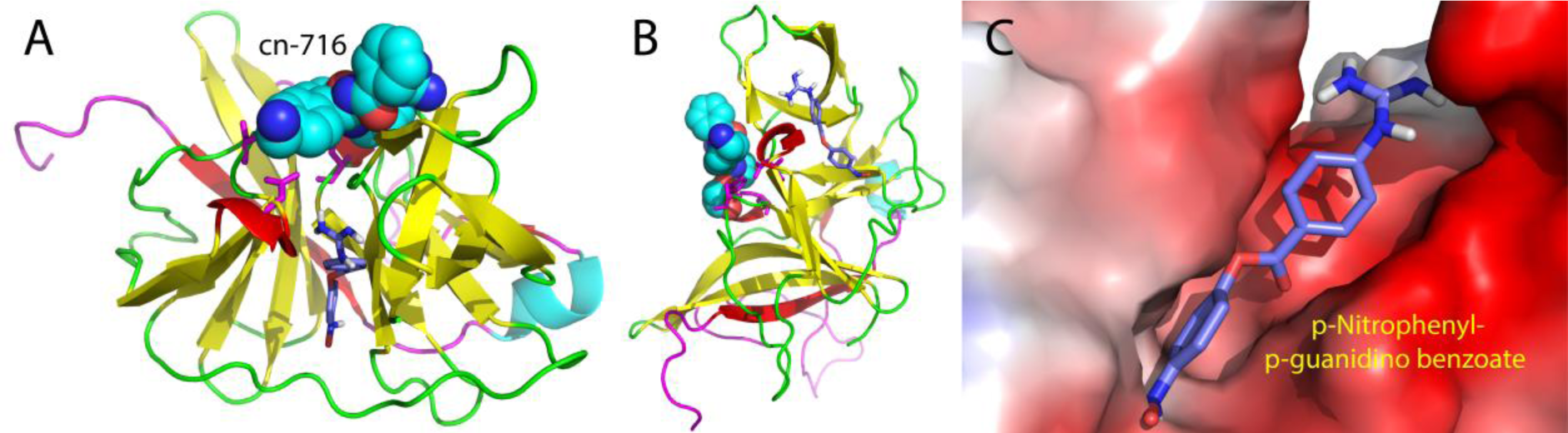
Binding pocket of Zika NS2B-NS3pro for p-Nitrophenyl-p-guanidino benzoate. (A)-(B) Structures of Zika NS2B-NS3pro with a peptidomimetic boronic acid inhibitor cn-716 in spheres and p-Nitrophenyl-p-guanidino benzoate in sticks. (C) The binding pocket in electrostatic potential surface for p-Nitrophenyl-p-guanidino benzoate.

Most strikingly, despite having very different structure scaffolds and inhibitory constants (Fig. 1). Qucertin also bind to the small pocket as pNGB (Fig. 3). This is further highlighted by Fig. 4.

**Fig 3.**
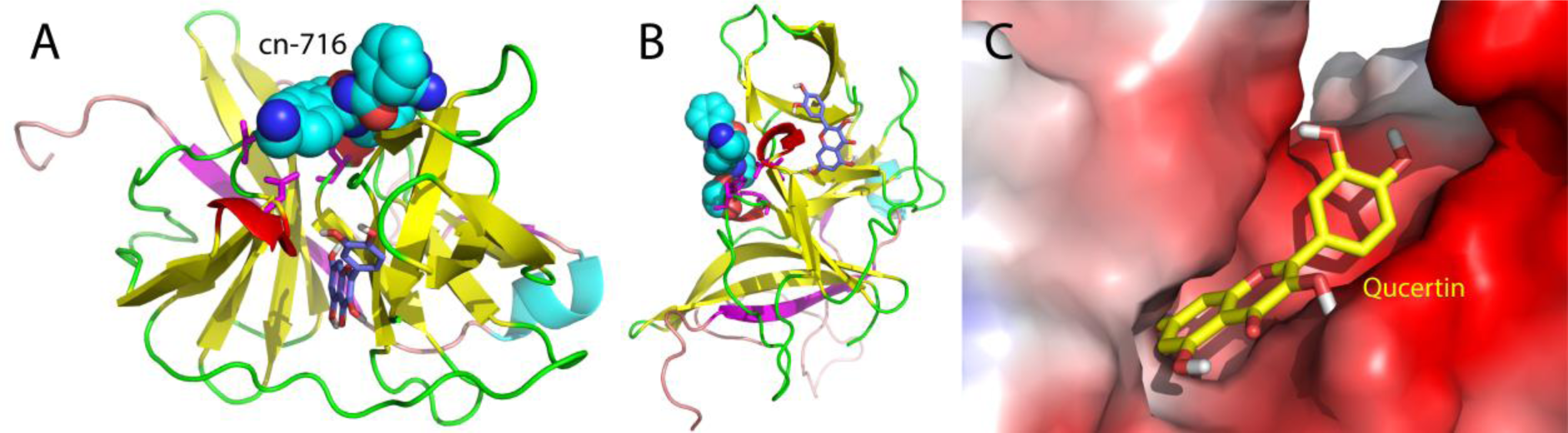
Binding pocket of Zika NS2B-NS3pro for Qucertin. (A)-(B) Structures of Zika NS2B-NS3pro with a peptidomimetic boronic acid inhibitor cn-716in spheres and qucertin in sticks. (C) The binding pocket in electrostatic potential surface for Qucertin.

**Fig 4.**
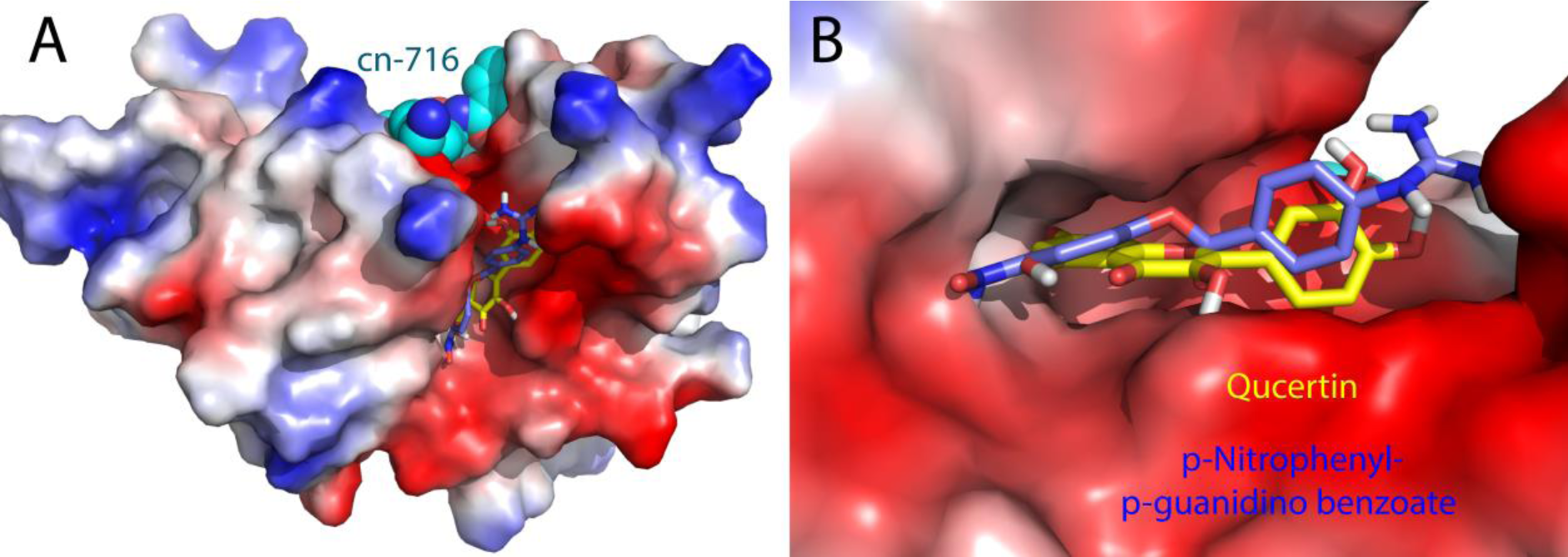
The same binding pocket of Zika NS2B-NS3pro for both p-Nitrophenyl-p-guanidino benzoate and Qucertin. (A)-(B) The binding pocket in electrostatic potential surface with both p-Nitrophenyl-p-guanidino benzoate and Qucertin in stickets, and cn-716 in spheres.

## DISCUSSION

Our study reveals that pNGB, a small molecule previously established as an active site inhibitor [19] is able to inhibit Zika NS2B-NS3pro at high affinity in non-competitive mode. This suggests that it acts as an allosteric inhibitor for Zika NS2B-NS3pro, fundamentally different from a recently reported peptidomimetic boronic acid inhibitor cn-716, which is a competitive inhibitor [25]. Indeed, molecular docking reveals that it binds to a pocket on the back of the substrate-binding pocket (Fig. 2). Interestingly, Qucertin with a different structure scaffold and binding affinity also binds to the same pocket. The indicates that this pocket of Zika NS2B-NS3pro is susceptible to allosteric inhibition, probably by many small molecules of different scaffols. Therefore, our discovery might open up a new avenue for the future development of allosteric inhibitors, which is highly demanded to achieve therapeutic inhibition of flaviviral NS2B-NS3pro complexes [25].

On the other hand, Quercetin is a flavonoid extensively found in many fruits, vegetables, leaves and grains, which have been used as an ingredient in supplements, beverages, or foods. It has been documented that quercetin has beneficial health effects ranging from antioxidant to nutraceutical [26]. Therefore, quercetin-containing fruits and vegetables may significantly benefit the public to immediately fight Zika virus.

Currently we are focused on optimizing various conditions which would allow atomic resolution studies on the structures, dynamics and inhibitor design for Zika NS2B-NS3pro by NMR spectroscopy [27–30].

## ACKNOWLEDGEMENT

This study is supported by Ministry of Education of Singapore (MOE) Tier 3 Grant R-154-002-580-112; Tier 2 Grant MOE2015-T2-1-111 and National Research Foundation of Singapore (NRF) R-154-002-529-281 to Jianxing Song. The funders had no role in study design, data collection and analysis, decision to publish, or preparation of the manuscript.

